# Functional and Structural Characterization of an IclR Family Transcription Factor for the Development of Dicarboxylic Acid Biosensors

**DOI:** 10.1101/2023.07.27.550818

**Authors:** Chester Pham, Mohamed Nasr, Tatiana Skarina, Rosa Di Leo, David H. Kwan, Vincent J.J. Martin, Peter J. Stogios, Radhakrishnan Mahadevan, Alexei Savchenko

## Abstract

Prokaryotic transcription factors (TFs) regulate gene expression in response to small molecules, thus representing promising candidates as versatile small molecule-detecting biosensors valuable for synthetic biology applications. The engineering of such biosensors requires thorough *in vitro* and *in vivo* characterization of TF ligand response as well as detailed molecular structure information. In this work we characterize the PcaR TF belonging to the IclR family. We present *in vitro* functional analysis of PcaR’s ligand profile and construction of genetic circuits for the characterization of PcaR as an *in vivo* biosensor in the model eukaryote *Saccharomyces cerevisiae*. We report the crystal structures of PcaR in the *apo* state and in complex with one of its ligands, succinate, which suggests the mechanism of dicarboxylic acid recognition by this TF. This work provides key structural and functional insights enabling the engineering of PcaR for dicarboxylic acid biosensors.

**Graphical Abstract:** 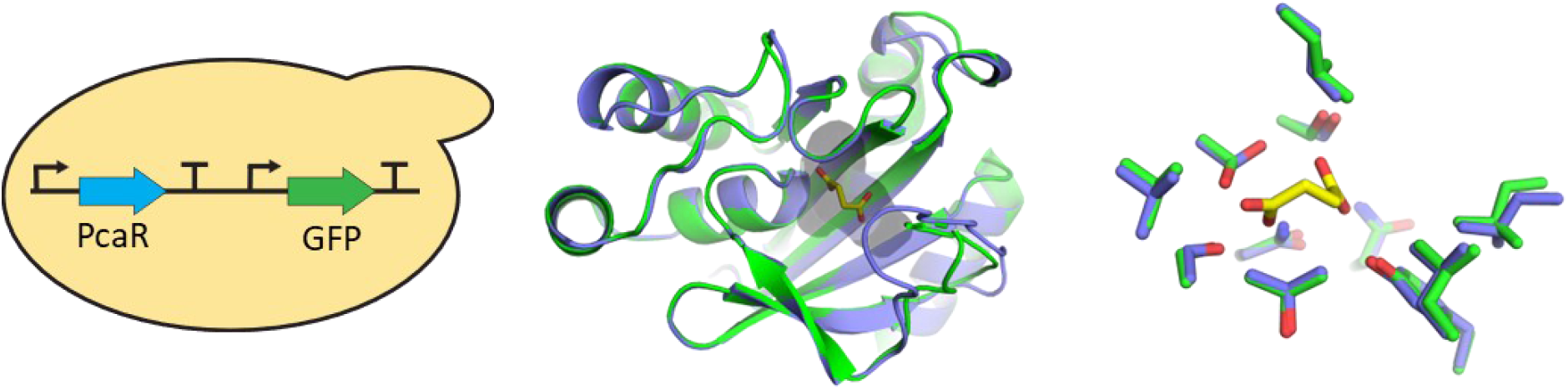

**Highlights:**

- PcaR is an IclR family transcription regulator responsive to dicarboxylic acids
- PcaR was established as an *in vivo* biosensor in yeast
- Crystal structure of PcaR in the *apo* form was solved
- Crystal structure with PcaR in complex with succinate was solved
- Sequence alignments unveil ligand-binding positions in the IclR family

## Introduction

Bacteria evolved complex genetic networks for gene regulation in response to a multitude of chemical cues [1]. Such regulation often involves specific transcription factors (TFs) that promote or inhibit the expression of gene clusters in response to changes in concentration of certain small molecules (their ligands). TFs alter their interaction with nucleic acids upon binding of appropriate ligands and this property has been leveraged for their implementation by bioengineers in a variety of artificial genetic circuits [2].

TFs are categorized into distinct protein families based on sequence and structural features, with the TetR and LysR families being the most widely distributed and known to regulate a variety of cellular functions [3]. Another notable TF family are the IclRs, which are involved in the regulation of diverse cellular processes such as primary and secondary metabolism, virulence, and quorum sensing [4]. The IclR family is a diverse TF family including members that function as activators, repressors, or with dual function, repressing some genes and activating others [4]. IclRs are generally characterized by a N-terminal helix-turn-helix (HTH) DNA binding domain (DBD) which dimerizes or tetramerizes to bind to target DNA, and a C-terminal ligand binding domain (LBD) which is connected to the HTH domain by an α-helix and loop. IclR family members can form dimers or tetramers when bound to the target promoter [4]. Generally, IclRs that act as repressors bind to the target DNA to occlude RNA polymerase (RNAP) complex binding, or inhibit the progression of the RNAP complex through the open complex [5]. In contrast, TF activators are involved in the recruitment of RNAP or access of the RNAP to the DNA for transcription. The specific activation mechanisms remain unclear since the preferred DNA locations for binding and activation remain unknown for IclR activators [5]. In addition, a small subset of IclRs have been reported to demonstrate both the repression and activation functions. An example of such dual role is described for PcaU, which binds to its cognate promoter as a repressor, but can also trigger the activation of the same promoter in the presence of its ligand protocatechuate. It has been suggested that changes in PcaU’s interaction with RNAP leads to activation at the promoter in the presence of the ligand, but no detailed analysis of such changes have been pursued [6]. Thus, much remains to be elucidated with respect to the IclR family’s basic mechanisms of DNA regulation.

Although there is no stringent consensus DNA sequence for the binding motif of the IclR family, two main types of motifs have been identified, both containing two-box binding sites [7]. However, some sub-groups of the family have been predicted to have three-box binding sites, indicating possible diversity in the oligomerization modes across family representatives [7]. The LBD of the IclR family consistently includes a conserved fold featuring the multiple-stranded, curved β-sheet lined on both sides by multiple α-helices [8]. Analysis of structurally characterized members of the IclR family suggested that deviations from this fold across this family are limited to amino acid sidechain positions within the binding pocket and to perturbations of the loop that is part of the binding pocket [9]. While the IclR family members appear to share the common architecture of DBD and LBD, variations in the relative orientation of the DBDs and LBDs is indicative of the flexibility of the linker connecting these domains and may contribute to the variety of functions and mechanisms demonstrated by IclR family representatives [10].

Overall, the IclR family contains TFs of a variety of functions and mechanisms. For example, TtgV from *P. putida* demonstrates broad ligand specificity towards aromatic compounds, suggesting that other TFs in the family may have similar profiles [5,11]. TtgV is a dimer in solution and binds to its target DNA as a tetramer, preventing RNAP from accessing the promoter region, making it a repressor [12]. Upon ligand binding, TtgV is released from its target DNA site, allowing transcription to occur [11]. In the case of the founding member of the family, IclR from *E. coli*, glyoxylate and pyruvate showed antagonistic effects [13]. Glyoxylate disrupted the TF-DNA complex *in vitro*, favoring the dimeric state, whereas pyruvate increased the binding of the TF to the promoter by stabilizing the binding in the active tetrameric state. Analysis of IclR from *T. maritima* suggests that binding to the ligand affects the oligomerization of the TF and its affinity for DNA, further linking tetramerization to ligand binding [14].

The diversity of chemical stimuli recognized by IclR family TFs makes them attractive synthetic biology tools for the design of biosensors responsive to small molecules. Many bacterial TFs have been repurposed as small molecule biosensors, where the presence of the target molecule in a given milieu could drive transcriptional activation or repression of a read- out/reporter gene [2]. Such biosensors have been useful for a variety of applications such as screening for environmental contaminants and point of care diagnostics [15,16], as well as metabolic engineering applications in which microbes are precisely engineered for heterologous production of industrially valuable molecules and chemicals [17].

Characterized IclR family TF-based biosensors include among others the protocatechuic acid-responsive PcaU and the 4-hydroxybenzoate-responsive PobR [18,19]. Thus, there is a vast untapped pool of potential biosensor circuits that could be constructed from this TF family. According to the manually curated GroovDB database, less than 20 members of the IclR family members are well-characterized with known inducers and components to construct biosensors, while the Pfam database identifies over 27,000 sequences belonging to the IclR family, highlighting the significant limitation in known inducers for the vast majority of IclR’s [20,21]. Furthermore, the sequence diversity across the IclR family is also indicative of the wide variety of mechanisms and molecules that IclR TFs can react to and that have yet to be identified.

Currently there are only five representatives of the IclR the family that have been structurally characterized in complex with their corresponding ligands, including: *E. coli* IclR in complex with pyruvate (PDB: 2O9A) or glyoxylate (PDB: 2O99) [13]; PobR with *para*- hydroxybenzoate (PDB: 5W1E) [22]; a PobR mutant with altered specificity bound to 3- hydroxybenzoate (PDB: 5HPI) [9]; and most recently pHbrR bound to 4-hydroxybenzoate (PDB: 7CUO) [23]. The structure of BaaR in complex with acetate in the binding pocket has also been determined (PDB: 5WHM), but this acetate molecule derives from the crystallization buffer [24]. Previous work has shown that BaaR maintains an open loop formation when binding to acetate in the binding pocket, unlike what is seen in other examples like pHbrR where binding leads to a closed loop, indicating that acetate is likely not a ligand of BaaR [23]. Such a low number of IclR-ligand complexes reflects the limited understanding of the basis of ligand recognition by IclRs and hinders rational design of IclR-based TFs as biosensors.

In the Gram-negative bacterium *P. putida*, the IclR family TF PcaR binds to a variety of promoters, regulates the degradation of aromatic compounds such as *p*-hydroxybenzoate, and in response to its inducer β-ketoadipate or its analog adipate, induces the expression of enzymes in the pca branch of the β-ketoadipate pathway encoded by the *pcaIJ* operon [25,26]. Previously, PcaR has been investigated *in vitro*, revealing that it forms a stable homodimer in solution, and it further oligomerizes into a dimer of dimers to activate the *pcaIJ* promoter by binding with its operator region overlapping with the -35 and -10 regions, which are regions critical for RNAP binding [27]. β-ketoadipate can increase the formation of the PcaR-RNAP-DNA complex, indicating transcriptional activation behavior by this TF [27]. PcaR was also found to negatively autoregulate its own promoter through binding to the region encompassing the -20 to the +4 positions [27]. This is like other members of the sub-group of atypical IclR family members that can act as both transcription activators and repressors [5].

Among the few representative IclR-based biosensors, PcaR has been previously constructed into a functional dicarboxylic acid responsive biosensor in *E. coli*, in which PcaR was engineered to regulate the expression of the TetA antiporter carrying the promoter of the *pcaIJ* operon in response to various dicarboxylic acids [28]. In published tests, PcaR responded to a range of dicarboxylic acids with 4 to 7 carbons (C4-C7), with the greatest specificity to adipic acid (hexanedioic acid) and pimelic acid (heptanedioic acid), but the lowest to succinic acid (butanedioic acid). The broad specificity of PcaR is in line with TFs from various families that have been classified as “generalists” [29,30]. The promiscuity of such TFs has been successfully exploited for engineering response to new ligands [31]. In the most recent example, PcaR was engineered towards a more specific response to adipic acid in its native *P. putida* using mutational analysis based on computational modeling of the protein’s 3D structure and docking of small molecule ligands [32].

Previous analysis of PcaR’s utility as a prokaryotic biosensor highlighted this TF’s plasticity of response to various dicarboxylic acids stimuli and suggests that the application of PcaR in biosensors can be further expanded. However, to our knowledge, all biosensor designs involving PcaR have so far only been adopted for prokaryotic cells, in line with this TF’s origin [28,32]. The implementation of PcaR as a biosensor in eukaryotic cells would demonstrate an orthogonal application of the PcaR circuit and expansion of its potential applications. Also, there has been limited exploration of the *in vitro* substrate recognition properties of PcaR, which could provide valuable tools for monitoring the engineering of this TF. Finally, elucidation of the experimental 3D structure of PcaR, particularly in an active complex bound to a ligand molecule, would provide new opportunities for more accurate structure-based engineering by providing details of the molecular basis for ligand recognition and conformational dynamics beyond what homology or *de novo* modeling tools like AlphaFold2 can provide [33].

In this study, we characterized PcaR through an *in vitro* thermal shift/differential scanning fluorimetry (DSF) assay and demonstrate an orthogonal application of PcaR as an *in vivo* biosensor in the model eukaryotic system *Saccharomyces cerevisiae* [27]. To characterize the ligand binding of PcaR, we tested a panel of saturated and unsaturated C4-C6 dicarboxylic acids that are of use in metabolic engineering and biocatalysis, including ligands that have not been tested as PcaR ligands in previous studies. In addition, we obtained structural insights into the PcaR ligand recognition mechanisms by solving PcaR crystal structures in the ligand-free (*apo*) and succinate-bound states. We compare PcaR to other homologous TFs in the IclR family, unveiling common binding characteristics that can inform future engineering of IclR family TF-based biosensors. Additional docking analysis further investigates the ligand specificity of PcaR. Taken together, this work facilitates the engineering of PcaR-based dicarboxylic acid biosensors, as well as their directed evolution toward novel acidic ligand specificities.

## Results

### In Vitro Ligand Specificity Screening of PcaR Reveals Preference for Succinate

Firstly, we pursued the characterization of the ligand specificity profile of PcaR *in vitro.* To explore this, PcaR was expressed in *E. coli*, purified, and tested with a range of dicarboxylic acids in the *in vitro* DSF assay [34]. In this assay, the PcaR thermal denaturation profile was most impacted by the presence of succinate when all compounds were individually evaluated at an equimolar concentration of 25 mM (Figure 1). This result contrasts with previous *in vivo* biosensor data where an *E. coli* strain expressing PcaR showed higher induction by adipate compared to succinate [35]. The melting temperature (T_m_), estimated by the point where relative intensity of the signal is 0.5, ranges from 49.6°C in the control sample containing no ligand, to 53.2°C for the sample containing succinate at 25 mM. Other compounds that caused significant although moderate increases to T_m_ included adipate, fumarate, glutarate, and malate, in decreasing order. This DSF data demonstrates the ability of PcaR to bind to a range of dicarboxylic acids, including malate and glutarate which have not been previously identified as ligands for this TF.

**Figure 1:**
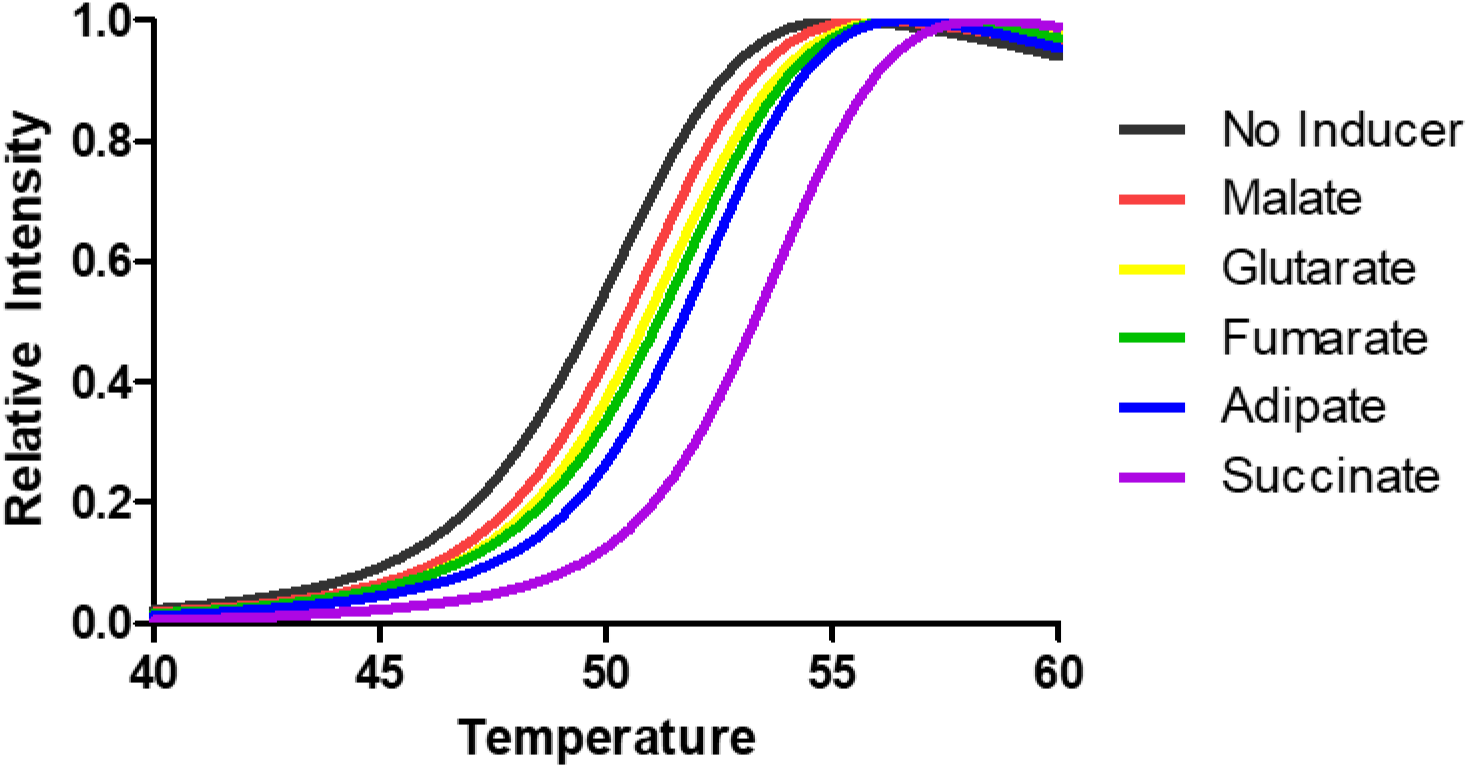
Differential scanning fluorimetry (DSF) assay data across different dicarboxylic acid inducers at 25 mM with n=3. Succinate binding produces the largest increase in thermal stabilization as reflected from the shifted melt curve, followed by adipate, fumarate, glutarate, and finally malate.

### Development of a PcaR-based Dicarboxylic Acid Biosensor in S. cerevisiae

To demonstrate the versatility of PcaR as a biosensor beyond previously developed prokaryotic systems, we investigated the activity of the TF in a model eukaryotic organism – the budding yeast *S. cerevisiae*. For this we incorporated the *pcaR* gene into a previously developed *S. cerevisiae* eukaryotic chassis and tested the ability of this strain to react to the molecules identified in our *in vitro* assay through induction of the expression of the green fluorescent protein (GFP) reporter gene [29,36]. Given the described activity of PcaR as a dual activator and repressor, we designed and constructed genetic circuits that were based on its repressor activity [5]. This required insertion of the PcaR-recognized operator sequence between the TATA box and the transcription start site for the GFP gene to sterically interfere with the binding or progression of RNAP [37]. With this strategy in mind, we built a chromosomally integrated PcaR genetic circuit in *S. cerevisiae*. In this circuit, we expressed *pcaR* from the strong constitutive promoter *P_TDH3_*. In the control strain we expressed the *envy* GFP variant using another strong constitutive promoter, *P_CCW12_* (Figure 2A) [38]. In the sensor strain we engineered a *P_CCW12_* promoter variant, which we called *P_CCW12O_* by introducing two consecutive copies of the *pcaO* operator sequence 11 bp downstream of the TATA box consensus sequence TATA(A/T)A(A/T)(A/G) localised from 133 to 125 bp upstream of transcription start site (Figure 2A) [39]. When the fluorescence outputs were measured by flow cytometry, the fluorescence showed a marked drop of ∼6-fold in mean population fluorescence values upon insertion of *pcaO* (Figure 2B). In the cultures containing increasing concentrations of the dicarboxylic acids identified in the DSF assay, the flow cytometric analyses showed an increase in fluorescence ranging from ∼1.5 fold in the case of adipic acid at 200 mM to ∼7 fold with fumaric acid at 25 mM (Figure 2C). Molecule concentrations were selected based on solubility, with the lower maximal concentration used in the case of the latter compound reflecting its much lower solubility. This data shows that PcaR can function as a transcriptional repressor in *S. cerevisiae*. PcaR is also shown to be responsive to different dicarboxylic acids in line with previously described PcaR-driven biosensors in prokaryotic cells [28,32].

**Figure 2:**
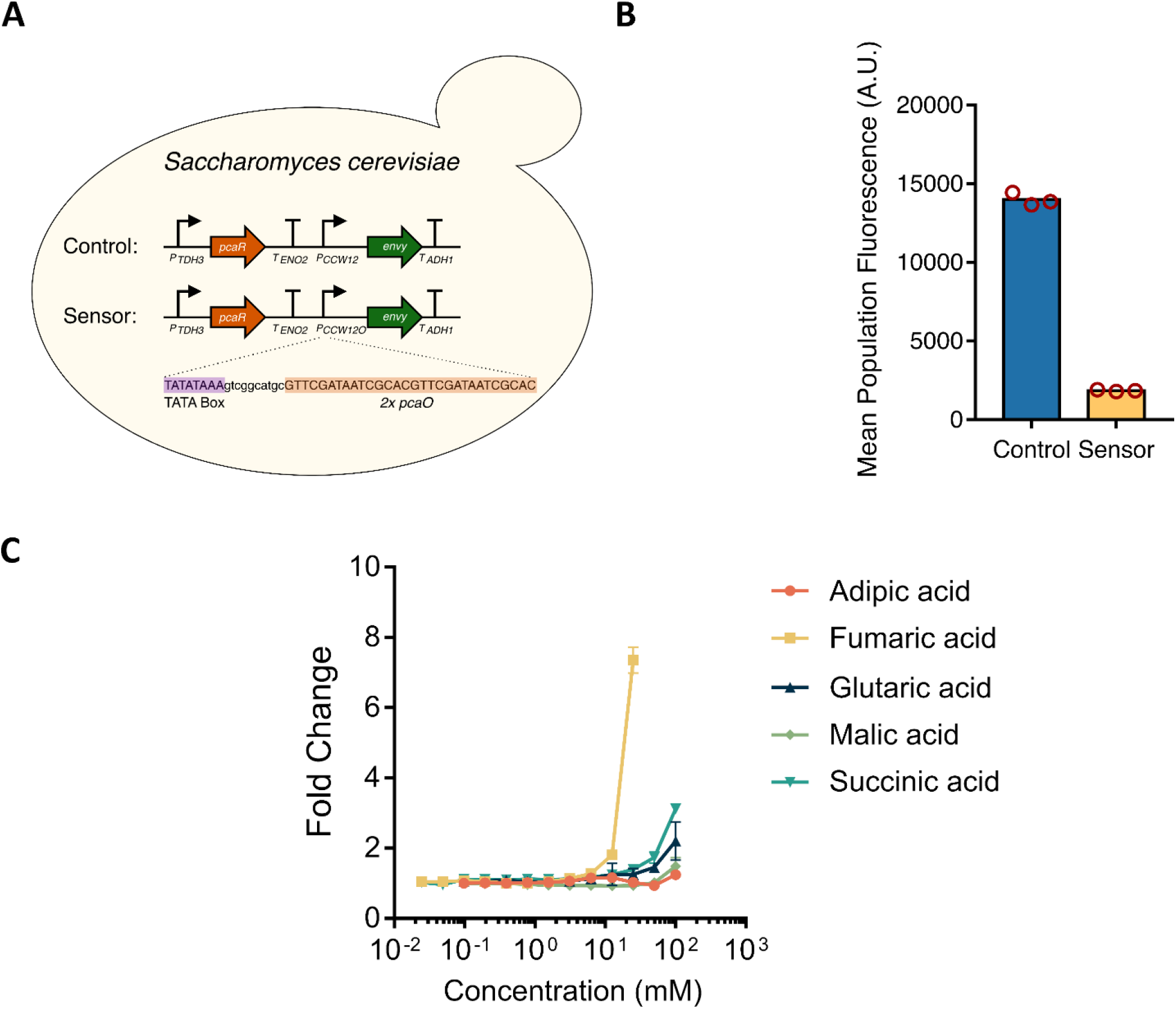
Construction and substrate profile of a *S. cerevisiae* PcaR biosensor. (A) Chromosomally integrated genetic circuit design in *S. cerevisiae*. PcaR is expressed from the strong constitutive promoter *P_TDH3_*. *P_CCW12_* is used for expressing *envy* GFP in the control strain, which was subsequently engineered in the sensor strain for dicarboxylic acid responsiveness by inserting two copies of *pcaRO* downstream of the terminal TATA box. (B) Mean population fluorescence of cells carrying the control and sensor circuit. (C) Changes in mean population fluorescence of the sensor strain in the presence of increasing concentrations of adipate, fumarate, glutarate, malate, and succinate.

### Crystal Structure of PcaR Reveals the Overall Fold Typical for IclR Family Members

To determine the molecular basis of ligand recognition by PcaR, we sought to structurally characterize the complexes between PcaR and ligands identified in our *in vitro* and *in vivo* assays. We determined the crystal structures of PcaR in the *apo* and succinate-bound states by the molecular replacement method using a previously determined structure from the homologous transcriptional regulator RHA06195 of *Rhodococcus sp.* RHA1 (PDB 2IA2) as a model (Figures 3 and 5; Table 1 contains data collection and refinement statistics). Like other previously characterized members of IclR family transcription regulators such as PobR [40], BaaR [41], BlcR [42], and pHbrR [23], the PcaR structure contains two polypeptide molecules in the asymmetric unit, reflecting the dimeric nature of this TF in solution. Following the IclR family’s characteristic architecture [5], each PcaR molecule consists of two domains, the N-terminal winged HTH DBD, and C-terminal LBD, with an α-helix linker region connecting these two domains (Figure 3A). The subunits are structurally similar and superimpose with an RMSD of 0.22 Å, forming an “X”-shaped homodimer crossed at the linker region. The loop between residues 154 to 169 that is localized to the binding pocket is not fully resolved in the observed electron density, indicating disorder. We assume that the movement of this loop could allow for modulating access to the main ligand binding cleft, as discussed later. There is a solute accessible opening to the binding pocket large enough to allow for entrance of a ligand at approximately 3.3 Å, however this opening likely fluctuates in size due to the movement of the abovementioned loop (Figures 3B and 3C).

**Figure 3:**
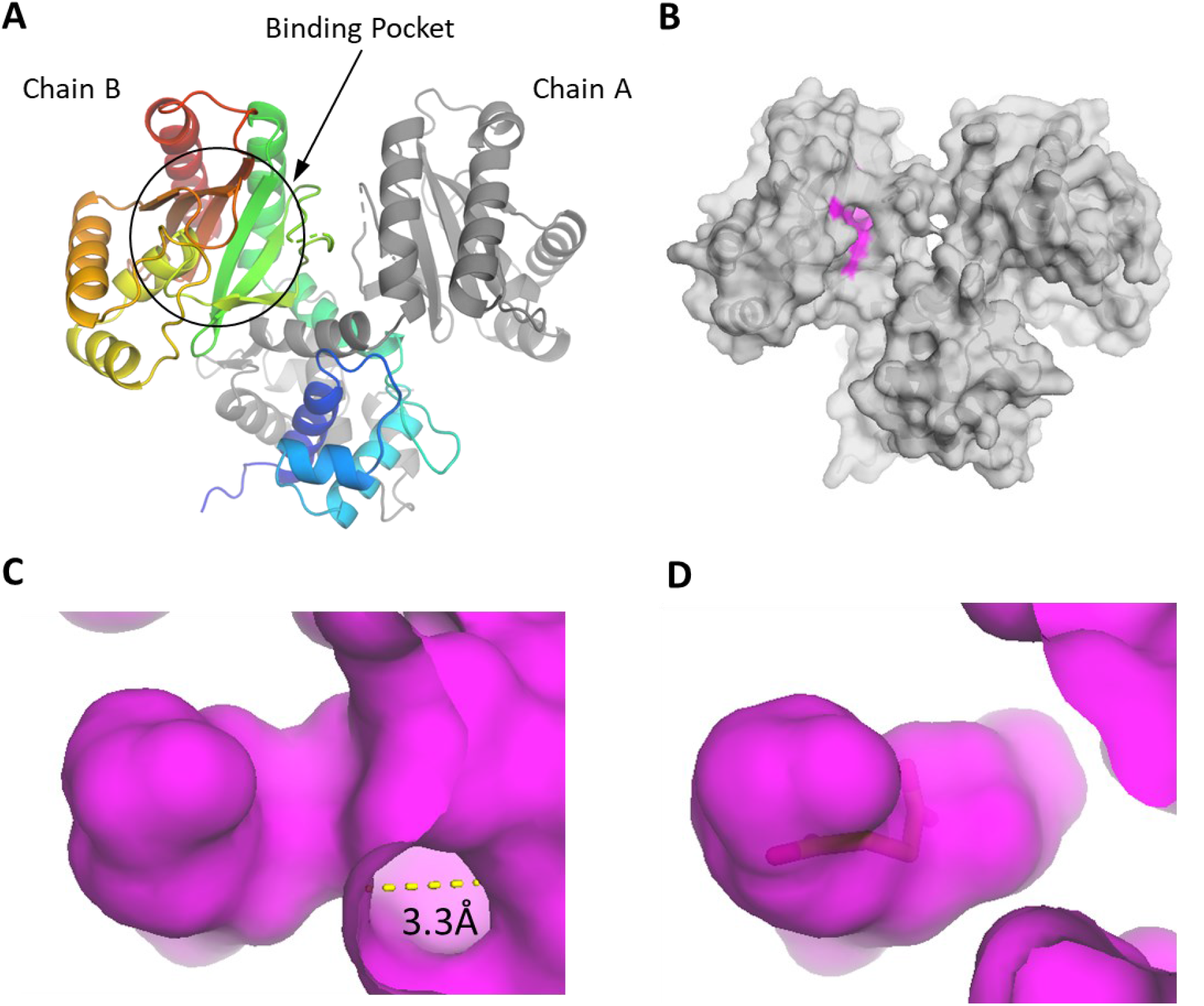
Overall *apo* structure of PcaR. (A) PcaR forms a homodimer in the asymmetric unit. The subunits contain an N-terminal winged helix-turn-helix DBD, α-helix linker, and C-terminal LBD. (B) Surface representation of the *apo* structure with interacting residues of the binding pocket highlighted in magenta. (C) Surface representation of the binding pocket in the *apo* structure. (D) Surface representation of the binding pocket when bound to succinate.

**Figure 4:**
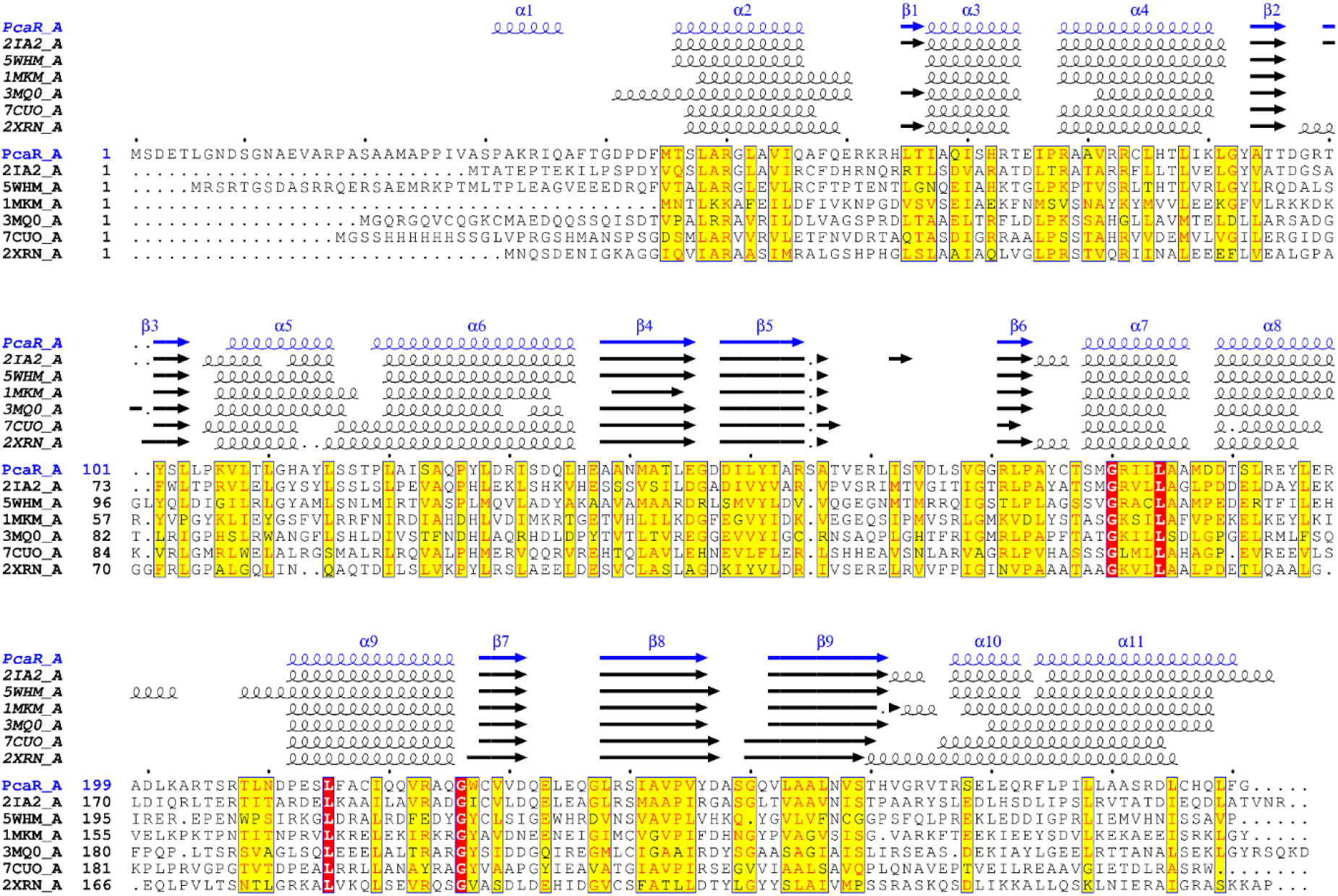
Sequence and secondary structure alignment of PcaR and homologues. The alignment indicates high conservation of secondary structure between PcaR and other members of the IclR family.

**Figure 5:**
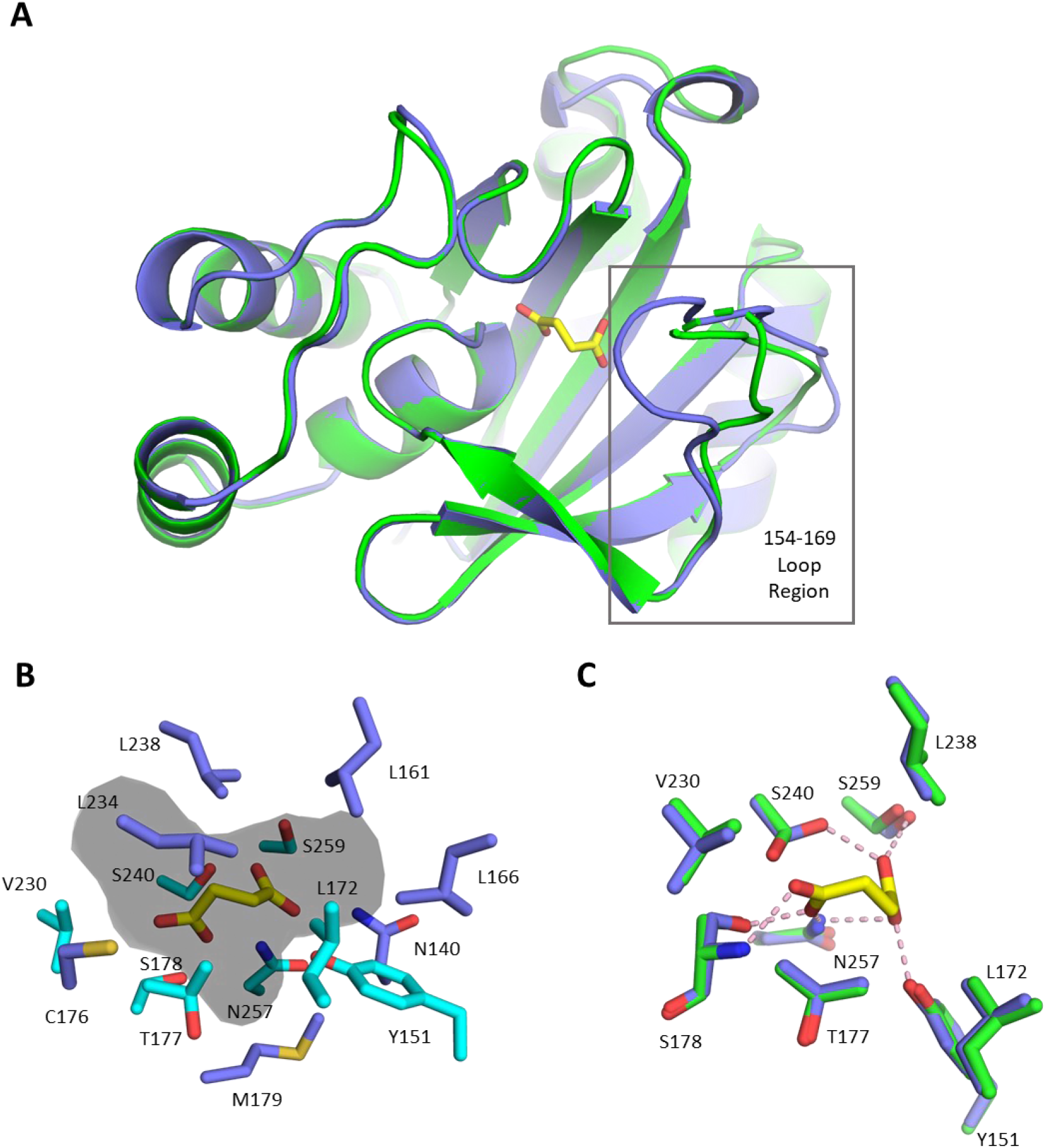
(A) View of the LBD with the ligand-bound (blue) structure overlayed on the apo structure (green) including succinate (yellow). A ligand-induced conformational change occurs at the loop region residues Arg154 to Gly169. (B) View of critical residues (cyan) and surrounding residues forming the binding pocket (dark colored) around succinate (yellow). (C) Apo structure (green) superimposed on ligand-bound structure (blue) focusing on the residues contributing interactions. Ligand binding altered the sidechain positions of some of the critical interacting residues and other residues in the binding pocket. Polar contacts are indicated (pink).

**Table 1:**
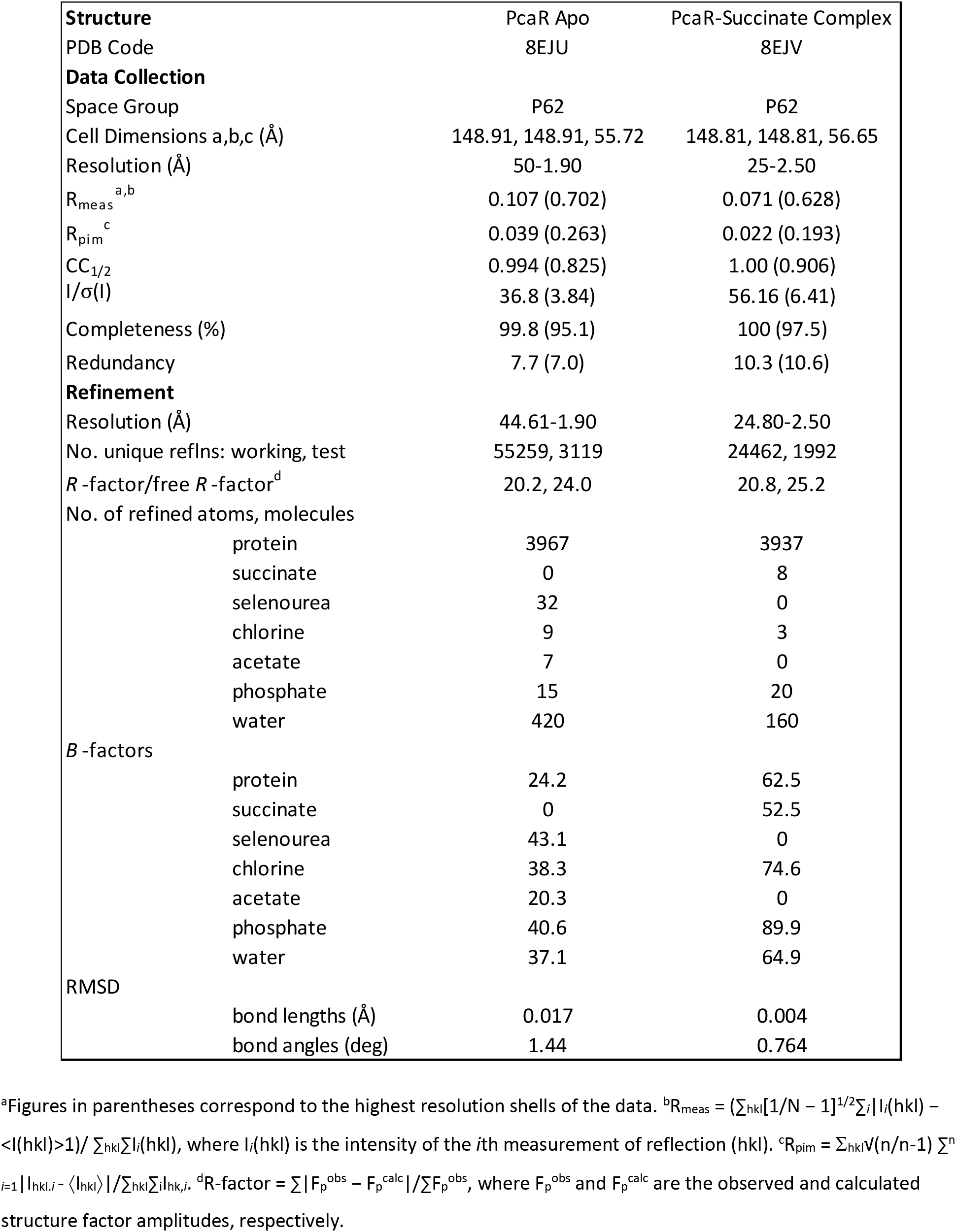
X-ray Diffraction Data Collection and Refinement Statistics.

To identify the family-wide conserved motifs across the IclR family, we conducted a structure-guided sequence alignment analysis using select structurally characterized IclR family members (Figure 4) [43]. The alignment shows high secondary structure congruence between PcaR and the other IclR family members. In particular, the residue corresponding to Pro77 has been shown to be highly conserved in previous analyses, suggesting its importance to DNA binding [5,7]. Out of the IclR family, only TtgV has been structurally characterized in complex with its operator sequence (PDB: 2XRO) [44]. In TtgV, Pro46 corresponds to Pro77 in PcaR and is placed within a sharp turn of the HTH domain and plays a role in the bending of the operator sequence upon DNA binding [45]. In addition, residues equivalent to Arg60 in PcaR also show high conservation in our alignment. In TtgV, this residue is shown to interact directly with the DNA, with variants of this residue being unable to bind to the DNA [45]. The DNA binding domain of TtgV and PcaR are structurally highly similar with an RMSD of 1.66 Å over backbone C-alpha atoms, suggesting that these TFs may share the same DNA binding mode.

### PcaR Undergoes Conformational Changes Upon Ligand Binding

We attempted to co-crystallize PcaR with the ligands identified in our experimental analysis. Out of such attempts, a succinate-bound complex structure was solved in the same space group as the *apo* structure, and the homodimer was again present in the asymmetric unit. This succinate-bound complex complements the *apo* structure as only the second pair of full-length crystal structures of an IclR family representative in both ligand-bound and unliganded form, with the only other one being from pHbrR [23]. Analysis of the ligand-bound structure reveals that PcaR accommodates succinate in a binding pocket of the C-terminal domain on both subunits of protein’s dimer (Figures 3D and 5A). The homodimer in the ligand-bound form retains an overall structure similar to that of the *apo* structure but undergoes key localized ligand-induced conformational changes. Similarly to the mechanism proposed for pHbrR, the exterior opening of the binding pocket in the PcaR/succinate complex closes upon the ligand’s binding (Figures 3C and 3D) [23]. Like other IclR family members including *E. coli* IclR, the binding pocket is centrally located and formed by six-stranded anti-parallel β-sheets and two long α-helixes (corresponding to α6 and α10 in PcaR) on one side and three (α7 to α9) short α-helixes on the other side (Figure 5A). These loops provide additional coverage to the ligand binding pocket, with the loop corresponding to residues Arg154 to Gly169 being the most extended. As seen in the PobR [9] and pHbrR complex structures with their respective ligands [23], there is a change in conformation of this PcaR loop upon binding to the ligand. While not completely resolved in the PcaR *apo* structure, Arg154-Gly169 loop appears to undergo a conformational change from an” open” to a “closed” formation more proximal to the binding pocket upon presence of the ligand, resulting in a decrease in the estimated binding pocket volume from ∼392 Å^3^ to ∼284 Å^3^. The binding of PcaR to succinate also affected the conformation of individual residues localized to the binding pocket (Figure 5C). For example, Ser240 undergoes a significant rotamer change upon ligand binding. However, we observed no other general conformational changes in the PcaR-ligand complex other than the change in the above-mentioned loop’s position, which was also the case for the pHbrR IclR TF [23]. Accordingly, we hypothesize that the change in loop position may play a key role in the mechanism by which ligand binding affects these TFs’ activities.

Our analysis of the PcaR-succinate complex structure identified several residues colocalized to the interior part of the binding pocket involved in interactions with this ligand (Figure 5C). Specifically, the sidechains of Tyr151, Ser178, Ser240, Asn257, and Ser259 form nine hydrogen bonds with the carboxylic groups of succinate. Critical residues lending interactions in the binding pocket belong to an interior binding pocket (Figure 5B) surrounding the ligand in the LBD in both subunits. Asn140, Tyr151, Leu172, Thr177, Ser178, Val230, Ser240, Asn257, and Ser259 also lend non-bonded contacts for positioning of the ligand in the binding pocket. Overall, the ligand binding mode of PcaR appears to be like other members of the IclR family, requiring only subtle conformational changes in the main binding pocket to enable ligand binding, but a larger conformational change and fixing in the position of the boundary loop 154-169.

### Structure-guided Sequence Analysis Elucidates Critical Residues for Ligand Binding Conserved Across the IclR Family

To compare the ligand binding site’s architecture in PcaR with other structurally characterized IclR TF’s, we aligned PcaR with the existing ligand-bound complexes with a focus on their LBDs (Figure 6). The hydrogen bonding network between the LBD residues and the ligand molecule appears to be limited in these TFs to a subset of residues lining the ligand binding pocket. In particular, the position corresponding to residue Ser178 in PcaR is occupied by residues involved in hydrogen bonding with the ligand in all structurally characterized ligand bound IclRs. In all analyzed IclR protein structures, this residue belongs to an α-helix inside the binding pocket and is typically occupied only by the small amino acids such as alanine, serine, and glycine. In PcaR, pHbrR, mutant PobR, and BaaR, the corresponding position is occupied by a serine, notable for its hydrophilicity and hydrogen bonding capacity. In *E. coli* IclR this position corresponds to a glycine, while in wild-type PobR this position is occupied by a hydrophobic alanine. In some of the IclR TF-ligand structures, the residue occupying the adjacent position on the helix is also involved in hydrogen bonding contacts with the corresponding ligand. PcaR structural analysis also reveals additional similarities in the ligand interaction residue network to IclR from *E. coli.* Specifically, Ser240 in PcaR corresponds to a cysteine in *E. coli* IclR, thus confirming the hydrophilic amino acid requirement for this position. This position falls on an interior β-sheet of the binding pocket. Similarly, the hydrophilic Asn257 in PcaR corresponds to a serine in *E. coli* IclR, while the BaaR structure also features an asparagine at this position. Finally, PcaR Ser259 on the interior β-sheet is also conserved in *E. coli* IclR.

**Figure 6:**
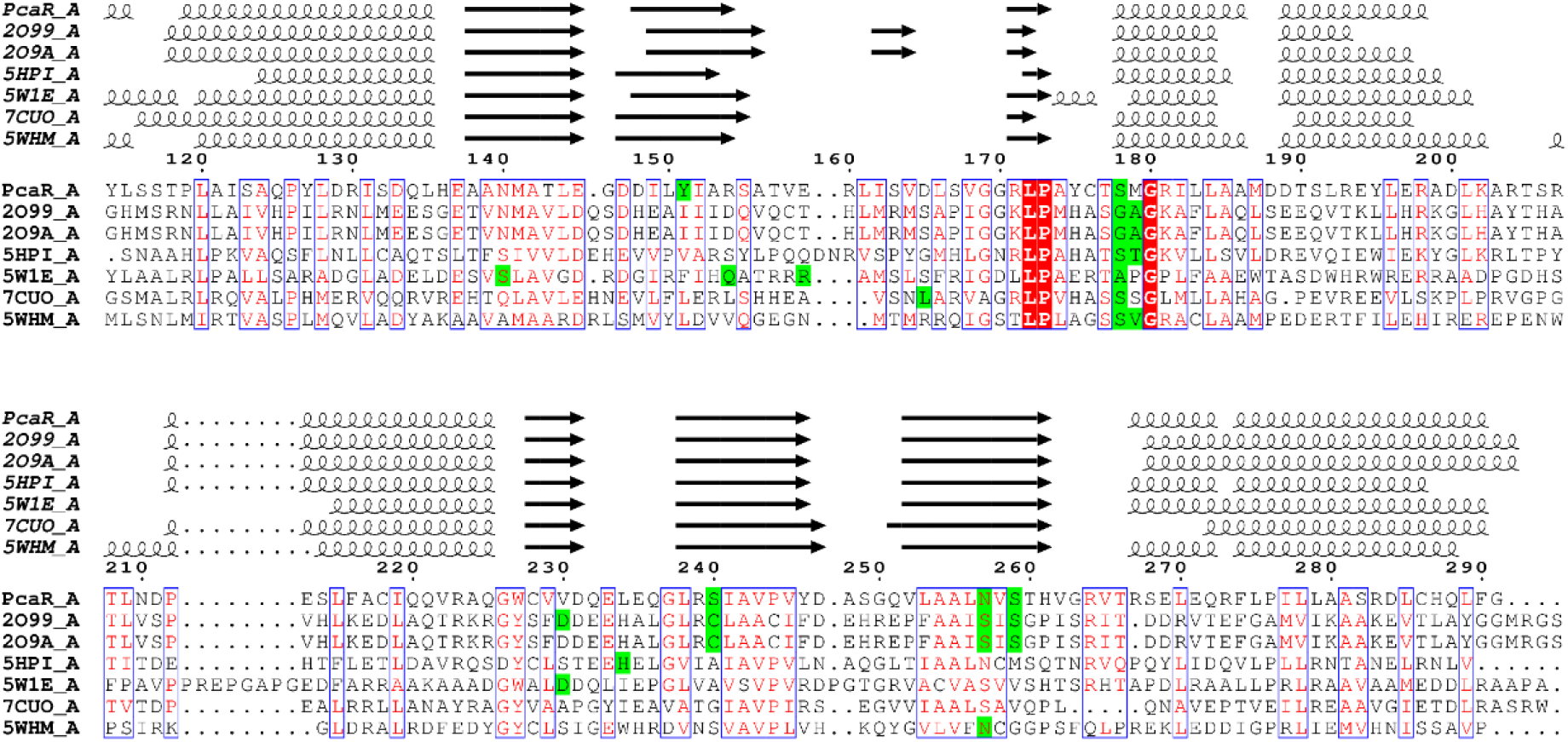
Sequence and secondary structure alignment of the LBD of PcaR and homologues with critical residues responsible for hydrogen bonding and the complexed molecule highlighted in green. Critical residues are homologous with some of the same positions responsible for hydrogen bonding with the complexed molecule in multiple proteins. In particular, the residue corresponding to 178 in PcaR is highly homologous and is involved in hydrogen bonding in all ligand-bound structures.

Overall, our analysis suggests that PcaR interactions with succinate follow the general pattern common among several IclR family members. The analysis reinforces the significance of gaining structural understanding of the specific binding pocket composition to allow for future exploitation by engineering recognition of novel ligands.

### Docking Studies Reveal Residues Involved in Ligand Discrimination in PcaR

Since our *in vitro* and *in vivo* analyses identified that PcaR can bind multiple dicarboxylic acid ligands, we were interested in investigating the rationale for such a ligand-binding profile based on details revealed by its complex with succinic acid, which we structurally characterized. To this end, docking studies were carried out with Rosetta and representative ligand binding models were analyzed in detail (Figure 7) [46]. The positions of residues in the binding loop where experimentally defined density was missing were modeled in fixed conformation, then PcaR was docked with the additional ligands identified in our above-described assays. While the ligand binding profiles are not identical, some notable trends can be observed. As seen in the alignment (Figure 6), all five ligands are bound to residue Ser178 through a hydrogen bond, reinforcing the importance of this residue in the binding pocket. Residue Thr177 contributes non-bonded contacts for binding of succinate and the other tested ligands. In the models, Ser240 interacts with all five of the ligands, forming hydrogen bonds with succinate, malate, and glutarate, while contributing non-bonded contacts towards adipate and fumarate. Ser259 also interacts with the five ligands, contributing a non-bonded contact to adipate, and hydrogen bonds to the other molecules.

**Figure 7:**
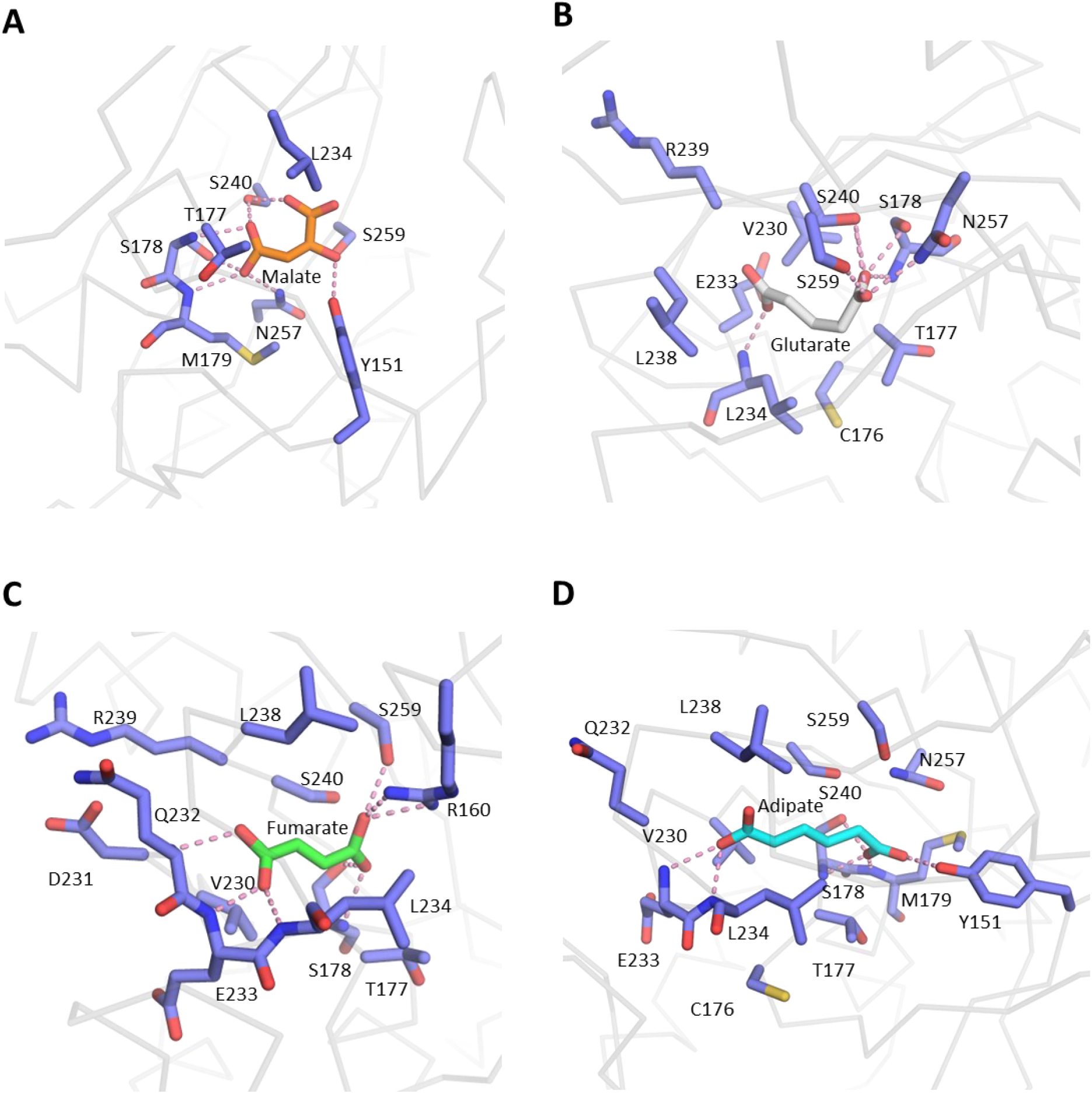
Predicted docked ligand complex models with PcaR and the tested dicarboxylic acid ligands. (A) Malate. (B) Glutarate. (C) Fumarate. (D) Adipate.

Next, we analyzed the docking models for possible rationale of PcaR’s small molecule effector specificity. Our analysis revealed two unique ligand-residue interactions. Firstly, only the succinate molecule interacts with PcaR’s Leu172 through non-bonded contacts. Secondly, fumarate forms exclusive interactions with Arg160 through direct hydrogen bonds. Such unique interactions for succinate and fumarate may be contributing to the specificity of PcaR towards these compounds. Our analysis also suggested that succinate and its oxidized form fumarate, both C4 dicarboxylic acids, are the only ligands capable of forming saturated hydrogen bond networks with the binding pocket residues. The modelling of complexes with the remaining molecules features fewer numbers of interactions, suggesting that their binding in the binding pocket may be less stable compared to that of fumarate and succinate. This observation is in line with the differences in the binding performance for these ligands observed in our functional assays. Malate, while also a C4 dicarboxylic acid, is a substituted dicarboxylic acid owing to an additional hydroxyl group. This alteration in structure may have contributed to the different fit in the binding pocket of PcaR.

Although this docking analysis does not fully account for the potential flexibility of the loop of the binding pocket, it does reinforce the importance of specific residues for ligand binding. This analysis suggests specific positions in the binding pocket to target for potential alteration of PcaR ligand specificity (i.e. those outside of Ser178/Thr177/Ser240).

## Discussion

In this work, we functionally and structurally characterized the PcaR TF from *P. putida*, unveiling new information on its specificity toward small molecule ligands and demonstrated its potential as an *in vivo* biosensor in yeast. This new data deepends our understanding of PcaR activity *in vitro* and *in vivo* and opens new opportunities to use this TF for engineering of biosensors in both prokaryotic and eukaryotic cells for synthetic biology and metabolic engineering applications.

Analysis of PcaR interactions with potential ligands by *in vitro* assays provided mechanistic details valuable for future biosensor engineering. We used the DSF assay that allows for monitoring of protein-small molecule interactions based on changes of the protein’s thermostability. Our data shows that among tested ligands, succinate showed the strongest effect on PcaR followed by adipate, fumarate, glutarate, and malate. This data did not match the pattern we observed for the *S. cerevisiae* strain where PcaR was engineered to repress GFP expression. In this *in vivo* set up, fumarate was the strongest inducer of GFP signal, followed by succinate, glutarate, malate, and adipate. Notably, both succinate and fumarate are C4 dicarboxylic acids. This suggests a preference of PcaR for binding saturated and unsaturated C4 dicarboxylic acids, over substituted C4 dicarboxylic acids like malate or the larger C5 and C6 dicarboxylic acids. Further docking analysis of the binding modes of the ligands suggested possible diversion in interactions in the binding pocket across this ligand set that can account for the observed differences in specificity. In particular, the unsaturated and saturated C4 dicarboxylic acids have all potential hydrogen bonds satisfied, while this is not the case for the substituted C4 or the larger C5 and C6 dicarboxylic acids. This highlights the importance of the shape and size of the binding pocket and the flexibility of the ligands.

Results from both the *in vitro* and *in vivo* characterization are also in contrast to previously published data where the PcaR-based biosensor in *E. coli* was reported to have the highest specificity towards adipate [28]. The variability of the specificity of the biosensor in different cellular contexts may be the result of differences in the construction of the genetic circuit, physiochemical and biochemical properties of the ligands such as membrane permeability, or intracellular conditions such as pH, on top of differences between organisms such as transport mechanisms or the metabolism of the tested compounds. Taken together, this discrepancy highlights the context dependence of biosensors as seen previously in the variable response of the same TF biosensor in *E. coli* and *S. cerevisiae* [29], and emphasizes the importance of taking multiple factors into account when transferring TFs between host organisms or when utilizing cell-free protein expression systems, as aspects in each system can have wide-ranging implications for the function of genetic circuits [15,47].

Our crystal structures of PcaR in the *apo* form and succinate-bound complex suggest that the conformational changes associated with ligand binding in the case of PcaR are limited to the loop region spanning residues 154 to 169, and changes in positions of specific residues in the binding pocket. A similar trend has been observed in other IclR family members. The analysis of the PcaR-ligand complex structure implicates specific residues involved in contacts between the inducer and the binding pocket. The comparative analysis suggests high structural conservation across the IclR family and indicates a convergence of positions and types of residues of the IclR LBD implicated in ligand binding interactions. In the previous engineering of PcaR towards higher specificity for adipic acid in *P. putida*, the highest performing variants featured mutations at the position 230 and 240, which our structural analysis identified as critical for interactions with the ligand [32]. PcaR Ser240 was found to directly bind to succinate through a hydrogen bond, highlighting the importance of this residue in ligand recognition. Additional studies would be required to determine the impact of each individual mutation on the modification of the specificity of PcaR. Such information regarding PcaR’s interactions with its ligand and its placement in the larger IclR family context can inform the design of new IclR- based biosensors and their engineering. Considering the rapid expansion of computational methods and algorithms, future studies could potentially use such ligand-binding information to predict TF specificity, expanding the repertoire of TFs to serve as biosensor chassis [48].

Our study confirms the role of PcaR as a general dicarboxylic acid sensing TF complementing the promiscuous or “generalist” TF toolbox, which already includes other generalist TFs such as CamR that is responsive to monoterpenes, and CmeR that is responsive to aromatics and indoles [29,30]. Generalist TFs are excellent candidates for directed evolution efforts to engineer biosensors to respond to new molecules due to their inherent promiscuity [49]. Our characterization of PcaR including the structural data presented here will inform future engineering of biosensors for the detection of dicarboxylic acids. In addition, our data paves the way for directed evolution and protein engineering efforts using PcaR to evolve TFs responsive to new molecules, ultimately enabling modulation of cellular activity in response to new chemistries in multiple host organisms.

## Summary

In this study we characterized the ability of *P. putida* PcaR to interact with an array of acidic ligands, establishing the expanded dicarboxylic acid specificity range for this TF. The expanded specificity range can be harnessed for future engineering and application of PcaR for detection of other similar molecules of interest. We also show the orthogonality of the PcaR biosensor system, demonstrating its activity in a eukaryotic organism for the first time. However, the specificity profile differences between our *in vitro* and *in vivo* evaluations, in addition to differences between our results compared to previous data reported for its activity in prokaryotic systems, illustrate the complexity in developing PcaR or any other TF-based biosensors. Structural characterization of PcaR with and without an inducer contributes the second known pair of crystal structures of a full-length IclR in its liganded and unliganded states, thus providing detailed insights into the structural mechanisms of the IclR family. In addition, we recognise that IclR family members share conserved positions within their binding pockets involved in ligand recognition. That information along with our docking analysis that suggests residues important for ligand specificity in PcaR, can inform residue targets for future specificity engineering in PcaR and the IclR family as a whole.

## Materials and Methods

### Protein Expression and Purification

*pcaR* (GenBank ID: L33795.1) was amplified from genomic DNA of *P. putida* and subcloned into the pMCSG53 expression vector, which codes for an N-terminal His6-tag and a TEV protease cleavage site. *pcaR* was expressed in *E. coli* BL21(DE3) Gold cells. LB media was inoculated with overnight cultures and cells were grown to an OD_600_ of 0.8 at 37 °C, chilled to 20 °C, and induced overnight with 1 mM isopropyl β-D-thiogalactopyranoside (IPTG). Cells were harvested via centrifugation at 4500 RPM, pellets were resuspended in binding buffer (50 mM Hepes (pH 7.5), 500 mM sodium chloride, 5 mM imidazole, and 5% glycerol (v/v)) and lysed by sonication. Insoluble cell debris were removed via centrifugation at 15000 rpm. The soluble cell lysate fraction was loaded on a Ni-NTA column pre-equilibrated with binding buffer, washed with washing buffer (50 mM Hepes pH 7.5, 500 mM NaCl, 35 mM imidazole, and 5% glycerol (v/v)), and the N-terminal His6-tagged protein was eluted with elution buffer (50 mM Hepes pH 7.5, 500 mM NaCl, 250 mM imidazole, and 5% glycerol (v/v)). Purified His6-tagged protein was then subjected to overnight TEV cleavage using 60 μg of TEV per mg of His6-tagged protein and dialyzed overnight against a dialysis buffer (300 mM KCl, 25mM Hepes pH 7.5, and 0.5mM TCEP). The His6-tag and TEV were removed by re-running the protein over the Ni-NTA column. The tag-free protein was then dialyzed against pre-crystallization buffer (10 mM Hepes pH 7.5, 300 mM KCl). PcaR product was identified by 12% SDS-PAGE. Protein concentrations were measured using the Bradford assay. The tag-free proteins were concentrated using a BioMax concentrator (MilliporeSigma, Toronto, ON, Canada) and passed through a 0.2 µm Ultrafree-MC centrifugal filter (MilliporeSigma, Toronto, ON, Canada).

### Thermal Shift for Ligand Binding

The thermal shift/differential scanning fluorimetry (DSF) assay is used to rapidly screen compounds that bind to PcaR through an increase in the protein melting temperature, indicating an increased stability upon ligand binding. The melting temperature is measured indirectly through the increase in the fluorescence when the reporter dye SYPRO Orange binds to hydrophobic parts of the protein as it unfolds. DSF experiments were performed on a BioRad CFX96 Real-Time PCR Detection System using the fluorescence resonance energy transfer channel. Samples of 25ul biological triplicates were prepared using 5 μg protein and ligand solubilized in the pre-crystallization buffer. Following a 45-minute incubation with the ligand, Sypro Orange was added to a 5x concentration and used as the reporter dye. Samples were further incubated with the dye for 15 minutes and centrifuged at 2000 x g for 30 seconds before starting the run. The cycling conditions included a 5-minute equilibration at 5 °C, after which the samples were heat denatured using a linear 5 to 95 °C gradient at a rate of 0.5 °C with 20-second intervals. Data shown in Figure 1 is the mean of the biological triplicates.

### S. cerevisiae Strains, Growth Conditions, and Transformations

*S. cerevisiae* manipulations were performed in the prototroph CEN.PK113-7D. Strains were grown at 30 °C for ∼16 or 48 hours in liquid or solid yeast extract-peptone-dextrose (YPD) medium, respectively. Hygromycin (200 µg/mL) and G418 (200 µg/mL) were added for plasmid selection as needed. Yeast cells were transformed by heat shock following the standard Gietz PEG/LiOAc protocol scaled down to a 25 μl volume [50]. Briefly, cells were incubated for 15 minutes at 30 °C, heat-shocked for 30 mins at 42 °C, and then followed by a recovery step in YPD for 16 hours. Cells were then plated on YPD plates containing the appropriate antibiotics for selection.

### S. cerevisiae Strain Construction

*pcaR* from *P. putida* was codon-optimized for yeast expression using the IDT tool and purchased from Twist Bioscience. Oligonucleotides were purchased from Thermo Scientific and suspended in nuclease-free water to a concentration of 100 μM. Sequences of *pcaR* and the oligonucleotides used are listed in Supplementary Table 1. *S. cerevisiae* strains were constructed using CRISPR-Cas9-mediated genomic integrations and *in vivo* DNA homologous recombination [51,52]. Yeast cells were transformed with a DNA pool that comprised: 1- PCR-amplified promoters, *pcaR*, *envy* GFP, and terminators with ∼40 bp homology to the preceding and/or following part, mixed in equimolar ratios (total DNA ∼1000 ng). 2- Two ∼500 bp homology arms to the genomic locus (200 ng each). 3- Linearized pCas-G418 and/or pCas-Hyg vectors encoding Cas9 (150 ng each) [53]. 4- gRNAs obtained by oligo-extension targeting the characterized chromosomal locus FgF16 (in which all integrations in this work were made) with 40 bp homology to the Cas9 plasmids to allow plasmid recircularization by *in vivo* homologous recombination (2 μL of unpurified PCR reaction per transformation) [54]. For constructing the control strain, the full *P_CCW12_* was amplified using primers MN527 and MN516. For the sensor strain with the operator insertion, *P_CCW12_* was divided into two parts with the operator acting as the homology arm in between them. Primer sets MN527 and MN664 and sets MN663 and MN516 were used to amplify both promoter parts. Transformations were recovered for 16 hours in YPD, after which they were plated on YPD plates with the appropriate selection (Hygromycin and G418 at 200 µg/mL each). Genomic integrations were confirmed by colony PCR and Sanger sequencing. Finally, the Cas9 plasmid was cured by streaking colonies two successive times on YPD plates with no selection. Sequences of promoters, gene expression cassettes, gRNA(s) and terminators are listed in Supplementary Table 1.

### S. cerevisiae Biosensor Characterization

Chemicals for PcaR *in vivo* biosensor characterization were purchased from Sigma Aldrich and dissolved in SC medium pre-adjusted to pH 4, to obtain stock solutions with the concentrations listed in parentheses: fumaric acid (25 mM), succinic acid (200 mM), adipic acid (200 mM), glutaric acid (200 mM), malic acid (200 mM).

Cells were grown overnight at 30 °C and 300 rpm in 96 deep-well plates (Greiner®) in an Infors HT shaker. The following day, different dicarboxylic acid solutions were serially diluted by 2-fold in SC medium pH 4 across a fresh 96 deep-well plate at a final volume of 500 μL per well. Yeast cells were then seeded into the wells using a Thermo Finipipette at a 1 in 100 dilution. Plates were incubated at 30 °C and 300 rpm in the same shaker for 20 hours, after which Envy GFP fluorescence was measured by flow cytometry using an Accuri C6 Cytometer (BD Biosciences).

Cells were diluted 1:5 in deionized water and measured at an average rate of ∼2000 events/second for a total of 10000 events. The mean fluorescence of the total ungated population was plotted for each molecule and/or concentration tested. Dose-response curve data points are averages of triplicates, shown either as three data points, or a single data point with an error bar representing standard deviation.

### X-ray Diffraction, Data Collection, Processing

Crystals were grown at room temperature using the vapor diffusion sitting drop method. For the *apo* structure crystal, the protein was in 0.3 M KCl, 10 mM Hepes pH 7.5, 0.5 mM TCEP, and the crystallization conditions were 1.8 M ammonium dihydrogen phosphate, 0.1 M sodium acetate pH 4.6, 1% (v/v) ethylene glycol, 2.5 (v/v) 1-butyl-2,3-dimethylimidazolium tetrafluoroborate, and soaked with selenourea. For the complexed crystal structure, the protein was mixed with 0.3 M KCl, 10 mM Hepes pH 7.5, 0.3 M NDSB201, with the reservoir solution containing 1.8 M ammonium dihydrogen phosphate, 0.1 M succinic acid pH 7, and 1.25% w/v benzyltriethylammonium chloride. Cryoprotection solutions included 7.5% glycerol.

Diffraction data at 100 K was collected at beamline 19-ID of the Structural Biology Center at the Advanced Photon Source, Argonne National Laboratory. Diffraction data was processed using HKL3000 [55]. The structures were solved by Molecular Replacement using Phenix.phaser [56] and an existing structure of homologous transcriptional regulator RHA06195 from Rhodococcus sp. RHA1 (PDB 2IA2). Model building and refinement were performed using Phenix.refine and Coot [57]. Atomic coordinates have been deposited in the Protein Data Bank with accession codes 8EJU for the apo structure and 8EJV for the succinate complex.

Protein structures were visualized in PyMOL [58]. Residues involved in interactions with succinate were identified using the PDBsum webserver [59]. Binding pocket volumes were estimated using Caver [60].

### Alignment

Selected sequences were aligned with ClustalW [61]. The alignment along with the PcaR structure was put forth for secondary structure alignment with the ESPript webserver which also pulls structures from the PDB [43].

### Docking

Missing residues in the *apo* PcaR structure were added with PyMOL and optimized by ModLoop [58,62,63]. Ligand conformers were generated using the BCL::CONF conformer ensemble generator [64]. The ligands were docked into the ligand binding pocket of PcaR using Rosetta with flexible sidechains [46]. 1000 models were generated and top ligand interface score models were selected as representative binding modes.

## CRediT authorship contribution statement

Chester Pham: Conceptualization, Formal analysis, Investigation, Methodology, Writing - original draft. Mohamed Nasr: Formal analysis, Investigation, Methodology, Writing - original draft. Tatiana Skarina: Investigation. Rosa Di Leo: Investigation. David H. Kwan: Supervision, Funding acquisition, Writing - review & editing. Vincent J.J. Martin: Supervision, Funding acquisition, Writing - review & editing. Peter J. Stogios: Investigation, Formal analysis, Writing - original draft, Investigation. Radhakrishnan Mahadevan: Supervision, Funding acquisition, Writing - review & editing. Alexei Savchenko: Supervision, Funding acquisition, Writing - review & editing.

## Abbreviations

TF: transcription factor
DSF: differential scanning fluorimetry
DBD: DNA binding domain
LBD: ligand binding domain
PDB: protein data bank
HTH: helix-turn-helix
DSF: differential scanning fluorimetry
RNAP: RNA polymerase

## Acknowledgements

We would like to thank Kemin Tan at the Structural Biology Center, Argonne National Laboratory, for x-ray diffraction data collection and processing.

## Supplementary Information

**Supplementary Table 1:**
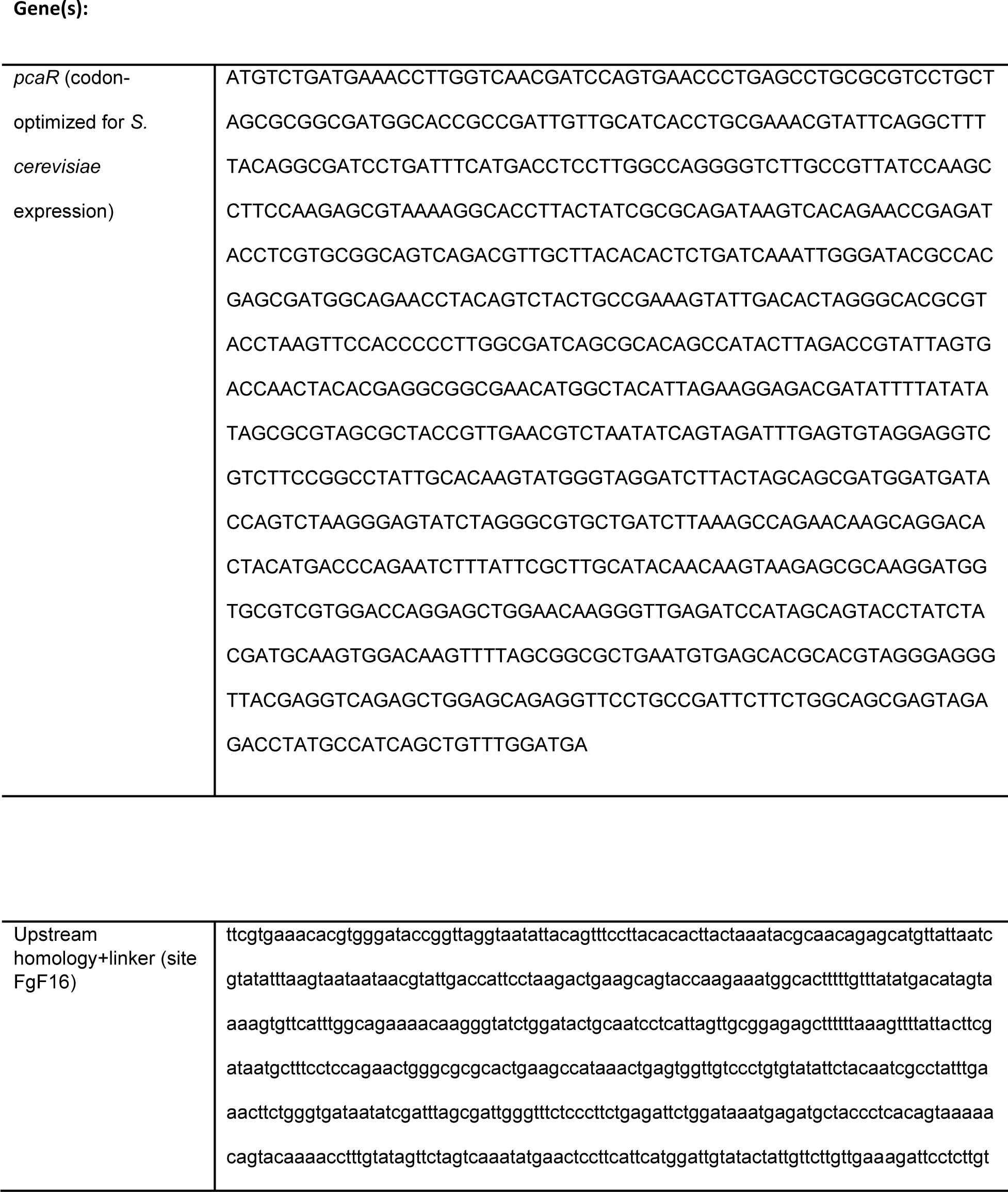
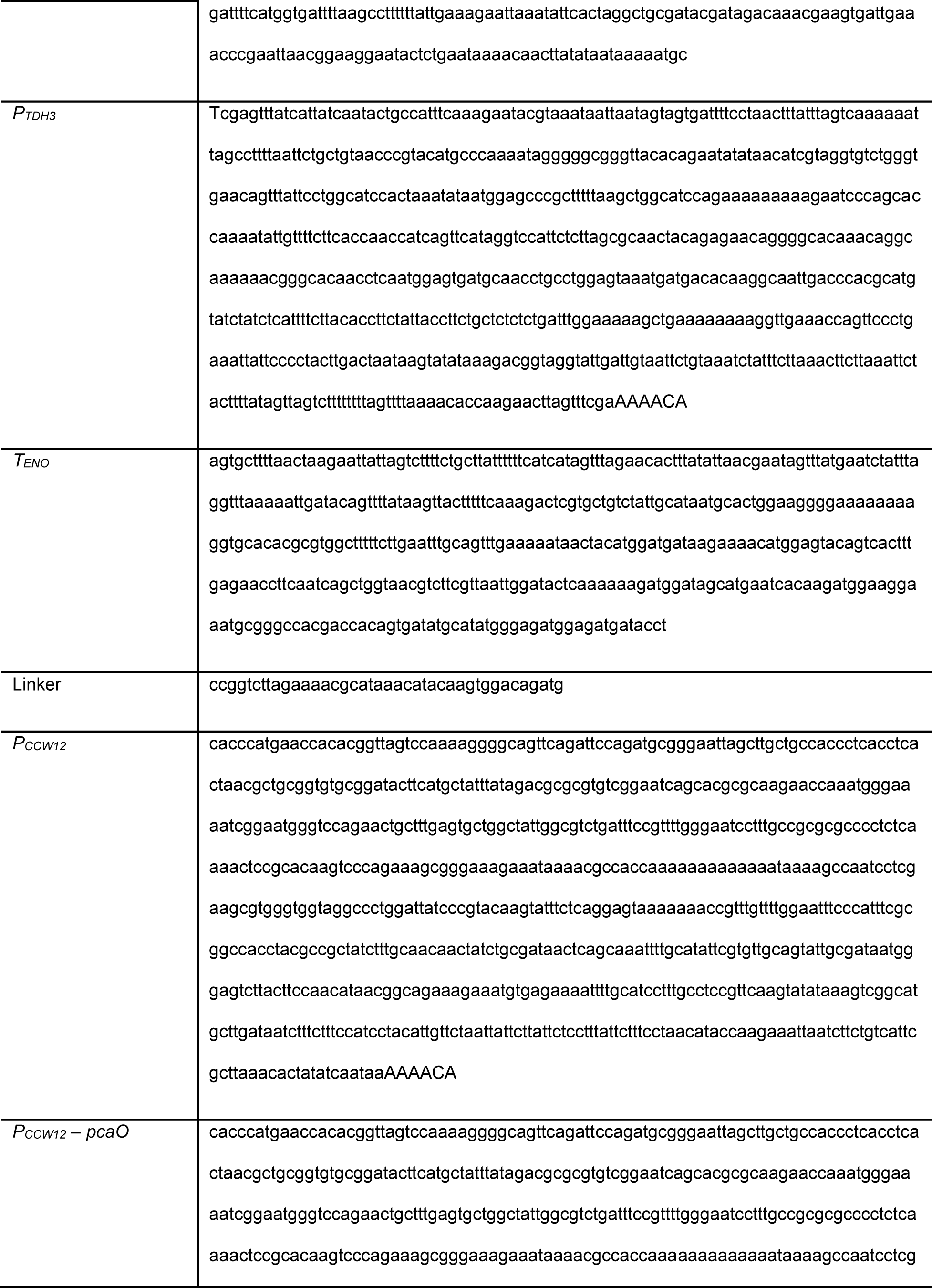
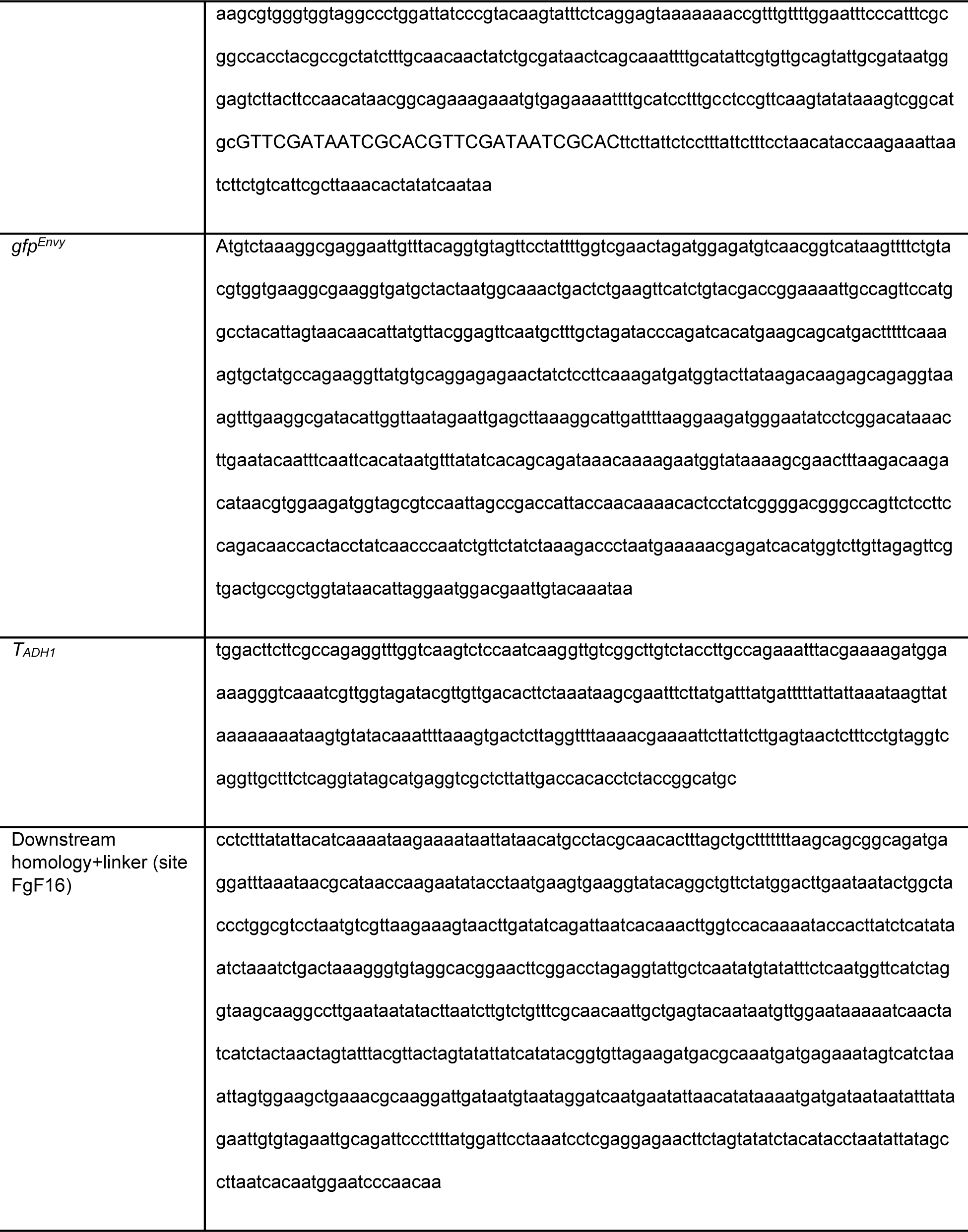
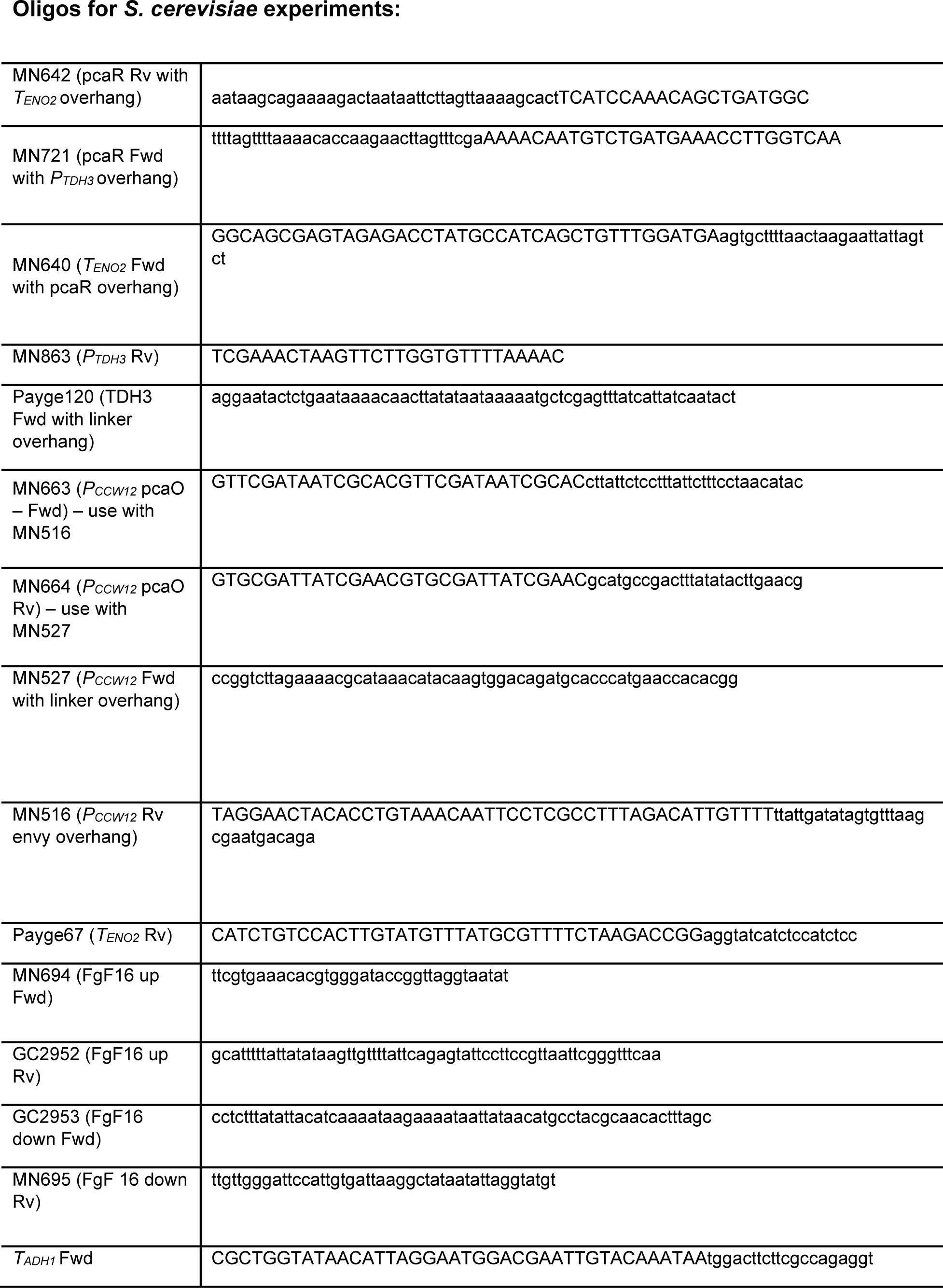
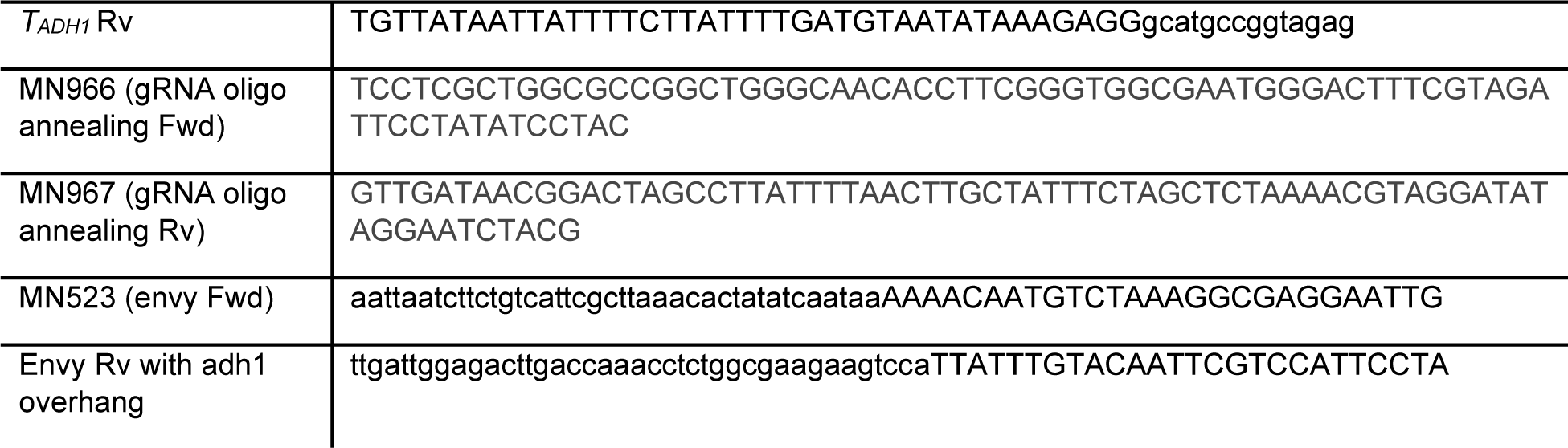

